# Not all pathways are the same – unique adaptations to submerged environments emerge from comparative seagrass genomics

**DOI:** 10.1101/2022.11.22.517588

**Authors:** Philipp E Bayer, Matthew W. Fraser, Belinda C. Martin, Jakob Petereit, Anita A. Severn-Ellis, Elizabeth A. Sinclair, Jacqueline Batley, Gary A. Kendrick, David Edwards

## Abstract

Seagrasses are an ecologically important group of plants that have returned to the sea from terrestrial ancestors on at least three occasions (Cymodoceaceae, Posidoniaceae, Zosteraceae). Seagrass-specific genomic adaptations to marine life are known from the Zosteraceae. However, independent lineages may have devised different solutions to life underwater. Here, we present two new genome assemblies from endemic Australian seagrasses, *Amphibolis antarctica* (Cymodoceaceae) and *Posidonia australis* (Posidoniaceae). We found large differences in genome size between *Amphibolis* and *Posidonia* driven by repeat expansion in *Posidonia*. We show that parts of ethylene pathways known to be lost in Zosteraceae are partially retained in older seagrass lineages (Cymodoceaeceae and Posidoniaceae). We describe adaptations within salinity, disease resistance, cell wall, and photosynthesis-related pathways not shared with other seagrasses. These findings provide insight into the impact of recolonising marine environments on formerly terrestrial plant genomes, with some adaptations previously thought to be universal to marine living not having occurred in *A. antarctica* and *P. australis*.

## Introduction

Seagrasses are a group of marine angiosperms in a unique ecological niche as the only flowering plants that can survive permanently submerged in saline waters. They exist in coastal environments surrounding every continent except Antarctica, forming critical habitats that contribute to ecosystem functions and services. Seagrasses support diverse food webs from microbes to megafauna, including commercial fisheries, improve water quality by increasing sediment settling rates and uptake of dissolved nutrients, and sequester vast amounts of carbon in underlying sediments (Aragones *et al*., 2006; Fourqurean *et al*., 2012; Fredley *et al*., 2019; Leigh *et al*., 2018). However, ecosystem functions and services continue to be eroded as global seagrass meadows decline. Historically, these erosions occurred due to poor water quality owing to adjacent anthropogenic land use. More recently, erosions have occurred due to climate-related impacts, such as marine heatwaves and extreme floods (Fraser *et al*., 2014). Marine restoration has been proposed as a tool to combat this global decline, but marine restoration has had mixed success at meaningful scales (Abelson *et al*., 2020; Fraschetti *et al*., 2021; Orth *et al*., 2020). More recently, the use of climate-resilient genotypes has been suggested to improve the success of marine restoration (Breed *et al*., 2019). Additional genome assemblies across seagrass genera will inform the conservation or restoration of these economically important species.

Seagrasses are polyphyletic as independent lineages have evolved separately from terrestrial ancestors in at least three return-to-sea events (Les *et al*., 1997). These events are reflected in three families: Zosteraceae, Hydrocharitaceae and Posidoniaceae/Cymodoceaceae within one order, Alismatales. These three families formed 100 MYA, with the ancestors of Zosteraceae and Cymodoceaceae diverging around 80 MYA (Janssen and Bremer, 2004).

The earliest study in this field used expressed sequence tags (ESTs) comparing expressed genes in *Zostera marina* and *Posidonia oceanica*, finding photosynthesis genes under positive selection (Wissler *et al*., 2011). This analysis was extended later using the first fully sequenced genome for a seagrass (*Z. marina*, Olsen *et al*. 2016), finding that the loss of ethylene and cell wall genes was associated with adaptation to a marine environment (Golicz *et al*., 2015). Similar findings were made in the *Z. muelleri* genome assembly (Lee *et al*., 2016). A recent study found ethylene genes EIN2, *CTR1*, and *ERS* present in Posidoniaceae and confirmed their loss in Zosteraceae (Chen *et al*., 2022). The latest Zosteraceae genome to be published was an improved version of the *Z. marina* assembly using long-read technology (Ma *et al*., 2021c). There is no published genome assembly for members of the Hydrocharitaceae, but genomic reads were used to investigate gene loss in Halophila ovalis (Hydrocharitaceae), finding that NADH dehydrogenase (*ndh*) genes are lost in *H. ovalis* but present in *Zostera* (Lee *et al*., 2018).

In this study, we assembled two new draft genomes from representatives of two Australian seagrass lineages that have independently returned to the sea, *Amphibolis antarctica* (Cymodoceaceae) and *Posidonia australis* (Posidoniaceae). These two species diverged shortly after the formation of Alismatales, approximately 95 MYA, while the *Amphibolis spp*. - Zosteraceae split was approximately 83 MYA (Kumar *et al*., 2017). This divergence represents a much earlier split among taxa for which whole genomes have been sequenced, as the ancestors of *Z. muelleri* and *Z. marina* diverged 39 to 21 MYA (Kato *et al*., 2003; Les *et al*., 2003). We compared the two new assemblies with the two published Zostera genomes (Zosteraceae; (Lee *et al*., 2016; Ma *et al*., 2021c; Olsen *et al*., 2016)) and other aquatic plants, such as duckweeds, algae and terrestrial plants. Our findings provide new insight into the genomic changes in plants associated with different pathways for adaptation to marine life.

## Results

### Two new genome assemblies for *Amphibolis* and *Posidonia*

We constructed genome assemblies of *Amphibolis antarctica* and *Posidonia australis* (Table 1, Supplementary Table 1) and compared the results with two existing genome assemblies for *Z. marina* and *Z. muelleri*. The two assembled genomes show a large difference in size, with an assembly size of 244 Mbp for *A. antarctica* and 1,215 Mbp for *P. australis*. These differences in assembly size are not reflected in the number of predicted genes, with 16,176 and 21,126 gene models for *A. antarctica* and *P. australis*, respectively (Supplementary Figure 1, 2, 3, 4, Supplementary Table 2, 3).

**Table 1.** Comparison of two published seagrass genome assemblies with two new seagrass genome assemblies

|  | *Z. marina* | *Z. muelleri* | *A. antarctica* | *P. australis* |
| --- | --- | --- | --- | --- |
| Reference | Ma *et al.* (2021c);<br>Olsen *et al.*<br>(2016) | Lee *et al.* (2016) | This study | This study |
| Estimated<br>genome size<br>(Mbp) | 259 | 889 | 233 | 1,399 |
| Genome<br>assembly size<br>(Mbp, excluding<br><1 Kbp contigs) | 202 | 632 | 244 | 1,215 |
| Number of<br>scaffolds > 1Kbp | 304 (6<br>pseudomolecules) | 35,334 | 16,916 | 258,843 |
| Number of<br>predicted gene<br>models | 20,450 | 35,875 | 16,176 | 21,126 |

### Convergent adaptations in gene content

We compared the gene content of *Arabidopsis thaliana* candidate genes linked with disease resistance, stomata development, ethylene synthesis and signalling, terpenoid biosynthesis, and chloroplast and mitochondrial genes across the four seagrasses (Figure 1, Supplementary Table 4, 5, 6). Annotation-based orthologues via Orthofinder (Emms and Kelly, 2019) and direct read-alignment of seagrass genomic reads with *A. thaliana* coding sequences (CDS) were compared for a complete overview.

**Figure 1.**
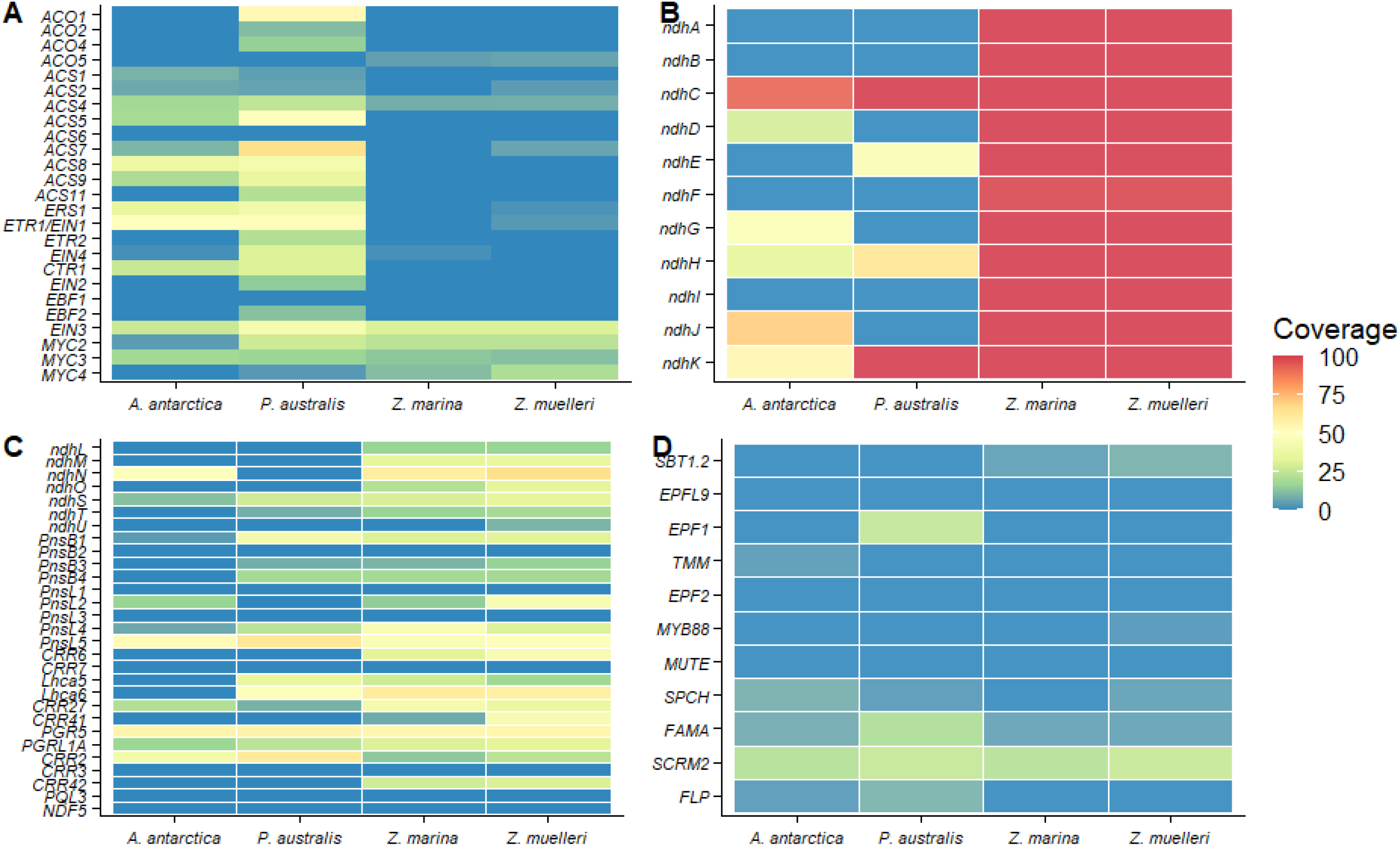
Gene coverage and gene size differences in the four seagrasses based on aligning sequencing reads of the five species with Arabidopsis coding sequences (A, C, D), or by comparing chloroplast gene annotation lengths with the Arabidopsis annotation (B). A) Ethylene pathway-related genes, B) ndh chloroplast genes, C) ndh-related nuclear genes, and D) stomata-related genes.

Seagrasses were found to contain very few disease resistance genes compared to terrestrial plants (Supplementary Figure 5). The genomes of *A. antarctica, P. australis, Z. marina*, and *Z. muelleri* contain only 739, 588, 755, and 1,256 NLR genes, respectively, which is much lower than in terrestrial plants such as *A. thaliana* (3,415 NLRs) or *B. distachyon* (3,058 NLRs). The RLP content is similar between terrestrial plants and seagrasses, with a median of 297 RLPs and 1,308 RLKs in the four seagrasses compared with 752 and 750 RLPs and 958 and 2,789 RLKs in *A. thaliana* and *B. distachyon*, respectively. Median RLK gene counts across seagrasses showed no reduction compared with terrestrial plants, with a median of 1,308 RLKs in seagrasses and 958 and 2,789 RLKs in *A. thaliana* and *B. distachyion*, respectively.

Ethylene genes were previously reported as lost in *Zostera* and *Halophila*, potentially due to the gaseous nature of the ethylene hormone and its lack of function in a submerged marine environment (Golicz *et al*., 2015; Lee *et al*., 2016; Olsen *et al*., 2016). In contrast, we found ethylene pathways are partially present in *A. antarctica* and *P. australis* (Figure 1A). The 1-aminocyclopropane-1-carboxylic acid (ACC) oxidases *ACO1, ACO2, ACO4*, and *ACO5* are lost in *A. antarctica, Z. marina* and *Z. muelleri*, but *ACO1, ACO2*, and *ACO4* are present in *P. australis*. Nine ACC synthases are lost in *Z. marina* and *Z. muelleri*, but all nine are present in *A. antarctica* and six are present in *P. australis*. The ethylene receptor *ERS1* is lost in *Zostera* but present in *A. antarctica* and *P. australis*. Seven ethylene receptors lost in *A. antarctica* (*CTR1, EBF1, EBF2, EIN1, EIN4, ETR2*, and *EIN2*) are present in *P. australis*.

*EIN3* was one of the few ethylene-pathway genes conserved in all seagrasses. We compared the *EIN3* proteins at the sequence level to determine whether these copies have diverged and perhaps neofunctionalised (Figure 2). The orthologues show a high level of divergence, except for a conserved region of approximately 150 amino acids at the beginning of *EIN3*. All terrestrial plant orthologues contain an uncharged region at 350 amino acids which is absent in three Zostera orthologues (*ZmuEIN3.1, ZmuEIN3.2, ZmaEIN3.1*). EIN3 orthologues exhibit size differences, ranging from 389 amino acids to 643 amino acids (median: 593 amino acids), yet all orthologues contain the ‘Ethylene insensitive 3’ Pfam domain (PF04873). These observations are consistent with a phylogeny of these sequences (Supplementary Figure 6), where seagrasses form a group distinct from terrestrial grass (*Oryza*), within which the *P. australis* orthologue *PaEIN3* shows the greatest sequence divergence from the Zosteraceae orthologues.

**Figure 2:**
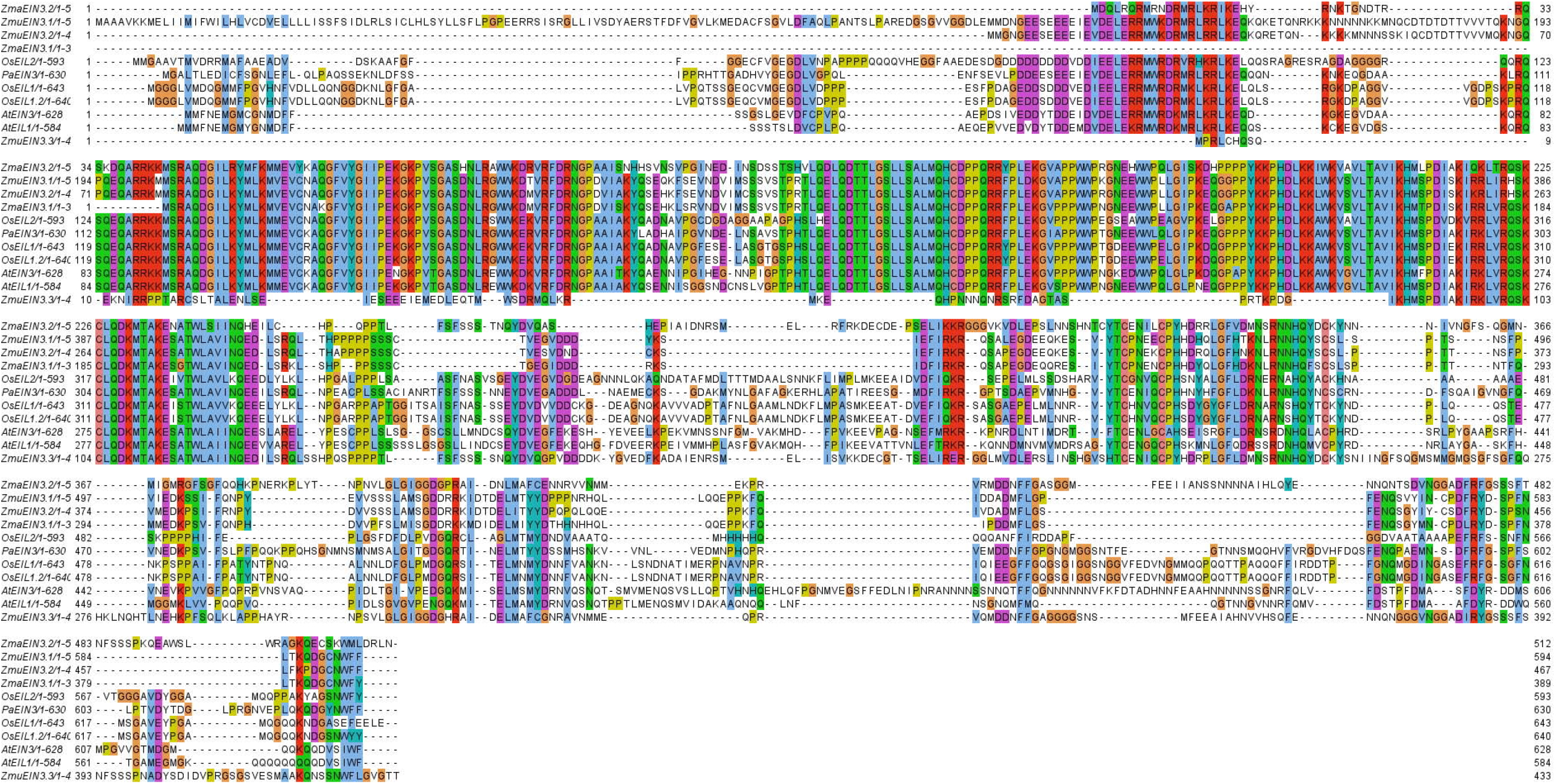
Multiple sequence alignment for the EIN3 orthogroup showing proteins of A. thaliana, O. sativa, and the three of the four seagrasses (no EIN3 orthologues in A. antarctica). Amino-acids are coloured by the ClustalX coluring scheme (orange: glycine G; gold: proline P; blue: small and hydrophobic amino acids A, V, L, I, M, F, W; green: hydroxyl and amine amino acids S, T, N, Q; magenta: negative-charged amino acids D, E; red: positive-charged amino acids R, K; dark-blue: histidine H and tyrosine Y).

To examine chloroplast-specific adaptations to marine life, we assembled and annotated the chloroplast genomes of *A. antarctica* and *P. australis* and reannotated the available chloroplast genomes of *Z. marina* and *Z. muelleri*. We then compared the chloroplast gene content with the *A. thaliana* chloroplast annotation to examine potential chloroplast and photosynthesis-specific adaptations to marine life. Chloroplast ribosomal genes show diverse presence and loss in the seagrass chloroplast genomes. The *rpl22* gene encoding chloroplast ribosomal protein CL22 was thought to be essential for survival in terrestrial plants, but deletion in *Nicotiana tabacum* resulted in no phenotypic changes (Fleischmann *et al*., 2011). *Rpl22* is not present in the plastomes of non-green plants *Eimeria tenella, Theileria parva*, and *Toxoplasma gondii* (Fleischmann *et al*., 2011) and legumes, where it was transferred to the nuclear genome (Gantt *et al*., 1991). In the seagrasses studied here, the *rpl22* gene is truncated by 17% to 24% in the *A. antarctica* and *P. australis* chloroplast annotations and absent in the *Z. marina* and *Z. muelleri* annotations, contrary to earlier studies in *Zostera* (Chen *et al*., 2021). Similarly, *rps16* is present only in the chloroplast genome annotations of *P. australis* and lost in the Zosteraceae.

Chloroplast *ndh* genes were previously thought central to photosynthesis but are lost in many orchids (Kim *et al*., 2015a; Lin *et al*., 2017), carnivorous plants (Lin *et al*., 2021; Wicke *et al*., 2014), and in some seagrasses, such as *H. ovalis* (Lee *et al*., 2018), but not *Z. marina* (Olsen *et al*., 2016). In the *A. antarctica* chloroplast annotation, only *ndhC* is present, while *ndhD, ndhG*, and *ndhH* are shorter by more than 50% compared to the *A. thaliana* genes, indicating they may have pseudogenised (Figure 1B, Supplementary Table 7). In *P. australis*, only *ndhC* and *ndhK* are completely present while *ndhE* is more than 50% shorter than the *A. thaliana* orthologue, indicating that it may have pseudogenised. The *ndh* complex is conserved and functional in *Zostera* (Lee *et al*., 2018; Ma *et al*., 2021b; Tan *et al*., 2020) which is confirmed by our *Z. marina* and *Z. muelleri* chloroplast re-annotations where all *ndh* genes are present.

Nuclear genes involved in *ndh*-complex-related pathways show a similar pattern (Figure 1C). Eighteen nuclear *ndh* genes are lost in *A. antarctica* and *P. australis* but conserved in *Z. marina* and *Z. muelleri*. Four genes are conserved across the four seagrasses: *PnsL5, PGR5, PRGL1A*, and *CRR2*.

Studies in *Z. marina* and *Z. muelleri* found genes involved in stomatal generation and differentiation to be lost (Golicz *et al*., 2015; Lee *et al*., 2016; Olsen *et al*., 2016). We found the same for *A. antarctica* and *P. australis* (Figure 1D). Only *SCRM2/SCREAM2*, a transcription factor involved in stomatal differentiation (Kanaoka *et al*., 2008), is preserved with one copy in *A. antarctica, Z. marina* and *Z. muelleri* and two copies in *P. australis*.

Salt Overly Sensitive pathways (SOS1-3) are the best-characterised salinity pathways in plants (Ji *et al*., 2013). Salinity-related GO-terms are enriched in genes present in all four seagrasses indicating a high degree of conservation. *SOS1, SOS2*, and *SOS3* are all present in seagrasses. *SOS2* has one orthologue in all seagrasses but two copies in *A. antarctica*. Only *SOS3* (also known as *CBL4*) shows several orthologues in the seagrasses, with one copy in *A. antarctica*, two in *P. australis*, three in *Z. marina*, and five copies in *Z. muelleri*. A phylogeny of these protein sequences showed considerable sequence diversity among seagrasses copies (Supplementary Figure 7). The phylogeny forms two groups, with one terrestrial group containing *O. sativa CBL4/SOS3* (*OsCBL4*) and multiple *Z. marina* and *Z. muelleri* copies, and a separate aquatic group. The additional *Zostera* copies *ZmaSOS3.2, ZmuSOS3.4*, and *ZmuSOS3.5* show considerable divergence not represented in the two included terrestrial plants.

A comparison of the *SOS3* orthologues at the protein-level (Figure 3) shows the core of the protein is highly conserved between *O. sativa* and *Z. marina* and *Z. muelleri*, while one *P. australis* copy shows substantial divergence with additional amino acids. All seagrasses contain additional uncharged amino acids at the beginning of proteins ranging from 13 amino acids in *Z. marina* and *Z. muelleri* to 205 amino acids in *A. antarctica*. We chose an additional set of 11 known *Arabidopsis* salinity resistance candidate genes (DeRose-Wilson and Gaut, 2011) and compared the presence/absence of these genes across seagrasses (Supplementary Table 5). Three to four salinity genes were lost, with *RDA29A* and *RDA29B* not present in *A. antarctica, P. australis, Z. marina*, and *Z. muelleri*, but present in the three duckweeds. The *A. thaliana* gene AT5G12010 involved in salt stress has no orthologue in *A. antarctica* or *P. australis*, but it has an orthologue in *Z. marina* and *Z. muelleri*.

**Figure 3:**
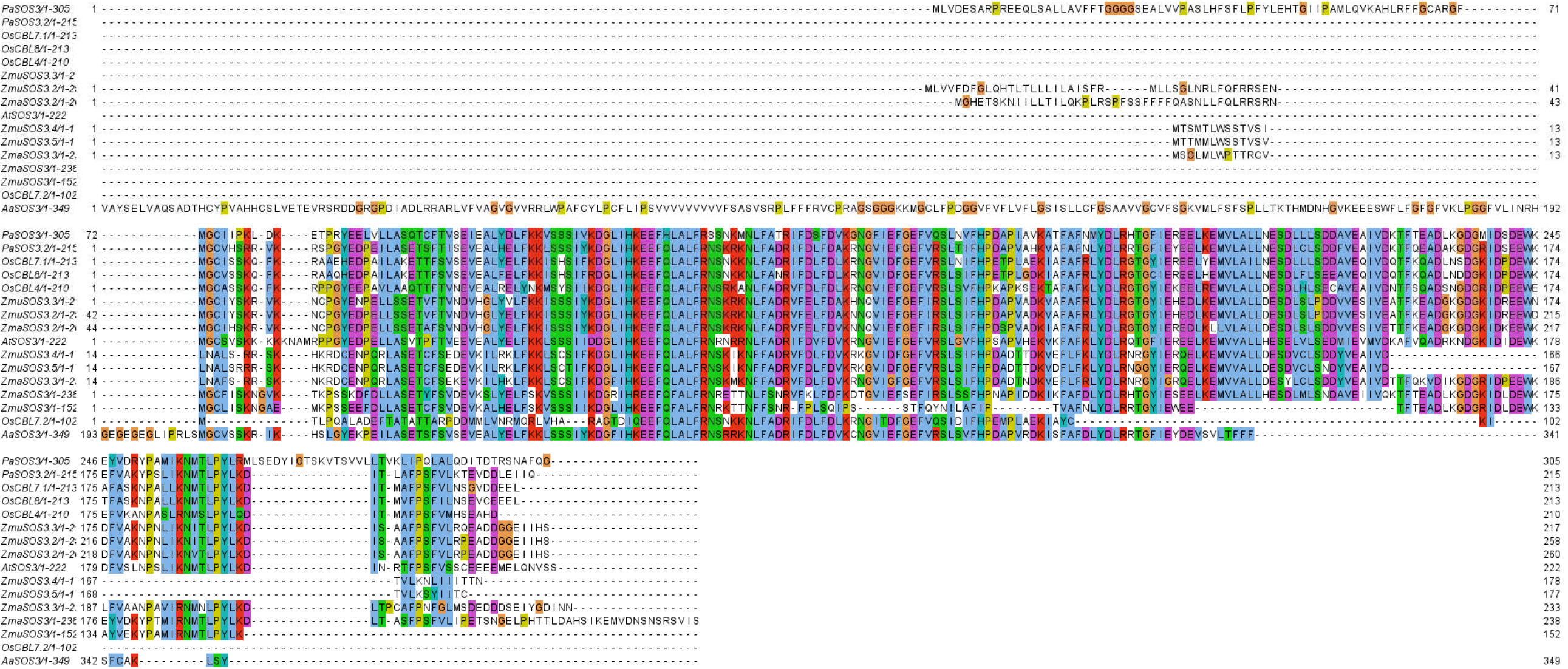
Multiple sequence alignment for the SOS3 orthogroup showing proteins of A. thaliana, O. sativa, and all four seagrasses. Amino acids are coloured using the ClustalX colouring scheme (orange: glycine G; gold: proline P; blue: small and hydrophobic amino acids A, V, L, I, M, F, W; green: hydroxyl and amine amino acids S, T, N, Q; magenta: negative-charged amino acids D, E; red: positive-charged amino acids R, K; dark-blue: histidine H and tyrosine Y)

*Z. marina* and *Z. muelleri* have lost all terpene biosynthesis genes (Lee *et al*., 2018; Olsen *et al*., 2016). *A. thaliana* terpenoid biosynthesis genes *TPS-CIN, TPS23, TPS03, TPS10*, and *TPS24* were clustered into one orthogroup containing no *A. antarctica* orthologues in this cluster, which is confirmed by no *A. antarctica* reads aligning with the *A. thaliana CDS*. The *P. australis* annotation, on the other hand, contains one terpenoid orthologue, as revealed by Orthofinder clustering. However, no *P. australis* reads align with the *A. thaliana* terpenoid biosynthesis genes, possibly due to sequence divergence.

We compared the gene content of the four seagrass genomes with the annotations of aquatic duckweeds (*Spirodela polyrhiza, Wolffia australiana*), microalgae (*Ostreococcus lucimarinus, Chlamydomonas reinhardtii*), and terrestrial plants (*Amborella trichopada, Arabidopsis thaliana, Brachipodium distachyon, Oryza sativa, Physcomitrella patens, Populus trichocarpa, Selaginella moellendorffii, Thellungiella salsuginea*,, *Vitis vinifera, and Zea mays*) by clustering protein-sequences using OrthoFinder. Duckweeds, microalgae, and seagrasses had a similar number of unique genes, with 1,498 (5%), 1,669 (5%), and 2,327 (8%) of orthogroups being specific to each respective group (Figure 4A, Supplementary Table 8). Most orthogroups included genes that are shared with terrestrial plants, with 6,058 (19%) orthogroups only containing genes from aquatic plants. Only 5,112 (17%) orthogroups had representative genes from all four groups, with orthogroups containing genes from all terrestrial plants alone, making up the bulk of orthogroups (13,748, 45%).

**Figure 4:**
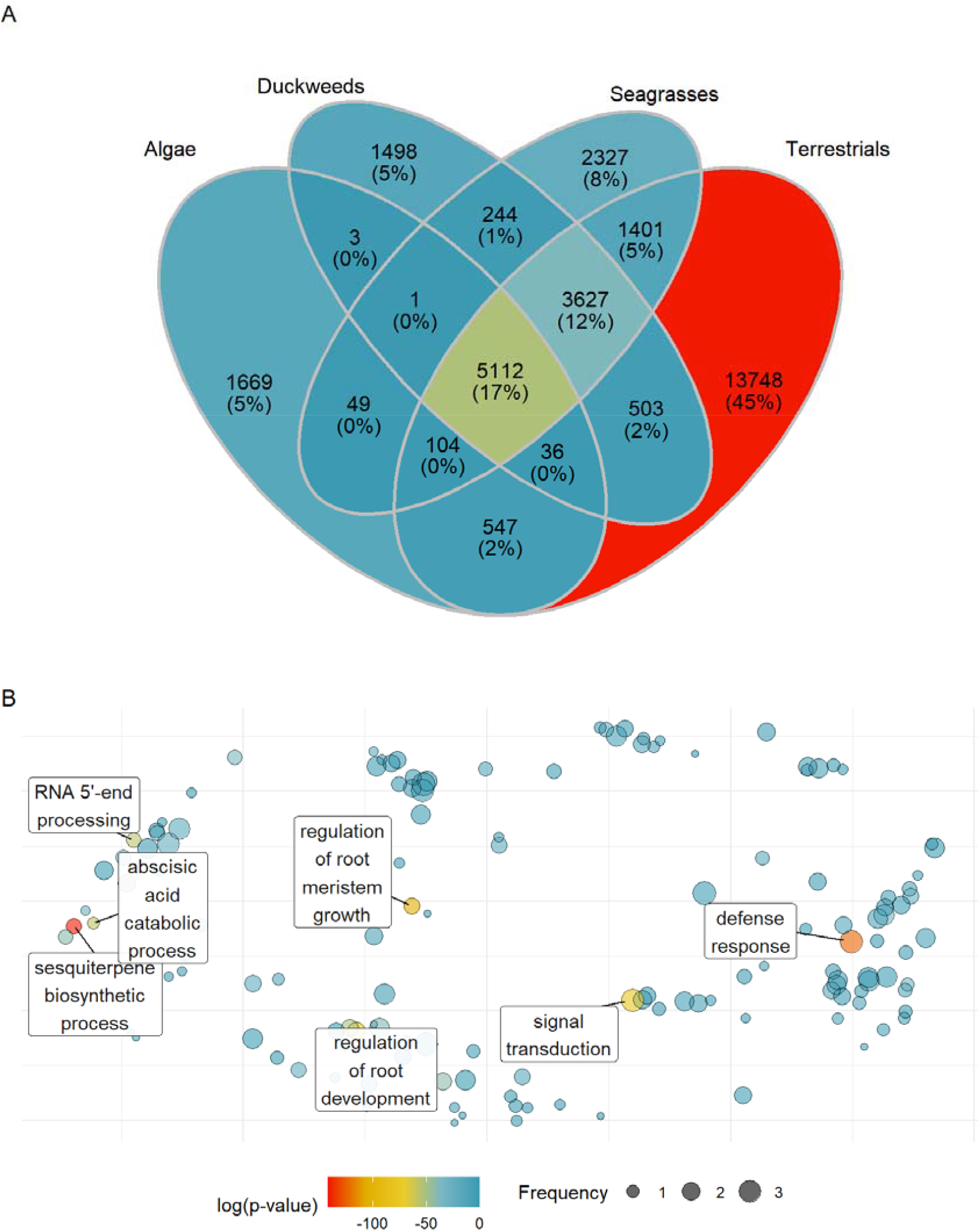
A) Shared and unique orthogroups between microalgae, duckweeds, seagrasses and terrestrial plants showing that just 17% of all orthogroups are shared across three groups, with the majority of unique orthogroups (45%) located in the terrestrial plants. B) Enriched Gene Ontology (GO)-terms in genes present only in terrestrial plants but lost in microalgae, duckweeds, and seagrasses. Presence of genes was determined by orthogroup membership. The GO-terms are grouped by semantic similarity using REVIGO (Supek et al., 2011). Size of terms is determined by how often similar terms appear in the GO-enrichment output. Dots are coloured by the log of GO-term enrichment adjusted p-values as determined by topGO, GO-terms with an adjusted p-value above 0.05 were removed (Alexa and Rahnenführer, 2009). The seven GO-terms with the lowest adjusted p-values are labelled.

We compared orthogroups unique and shared across the four seagrasses to examine seagrass-specific adaptations to underwater life (Figure 5A). The majority (6,534, 51%) of orthogroups were present in all four species. Of these, 2,983 (24%) were present in both *Zostera* species, but not in *A. antarctica* and *P. australis. Z. muelleri* had the largest number of unique orthogroups, with 1,262 (10%), consistent with its known whole-genome duplication event and subsequent functional diversification. Few orthogroups were present only in the two seagrasses *A. antarctica* and *P. australis* (1,421, 11%).

**Figure 5:**
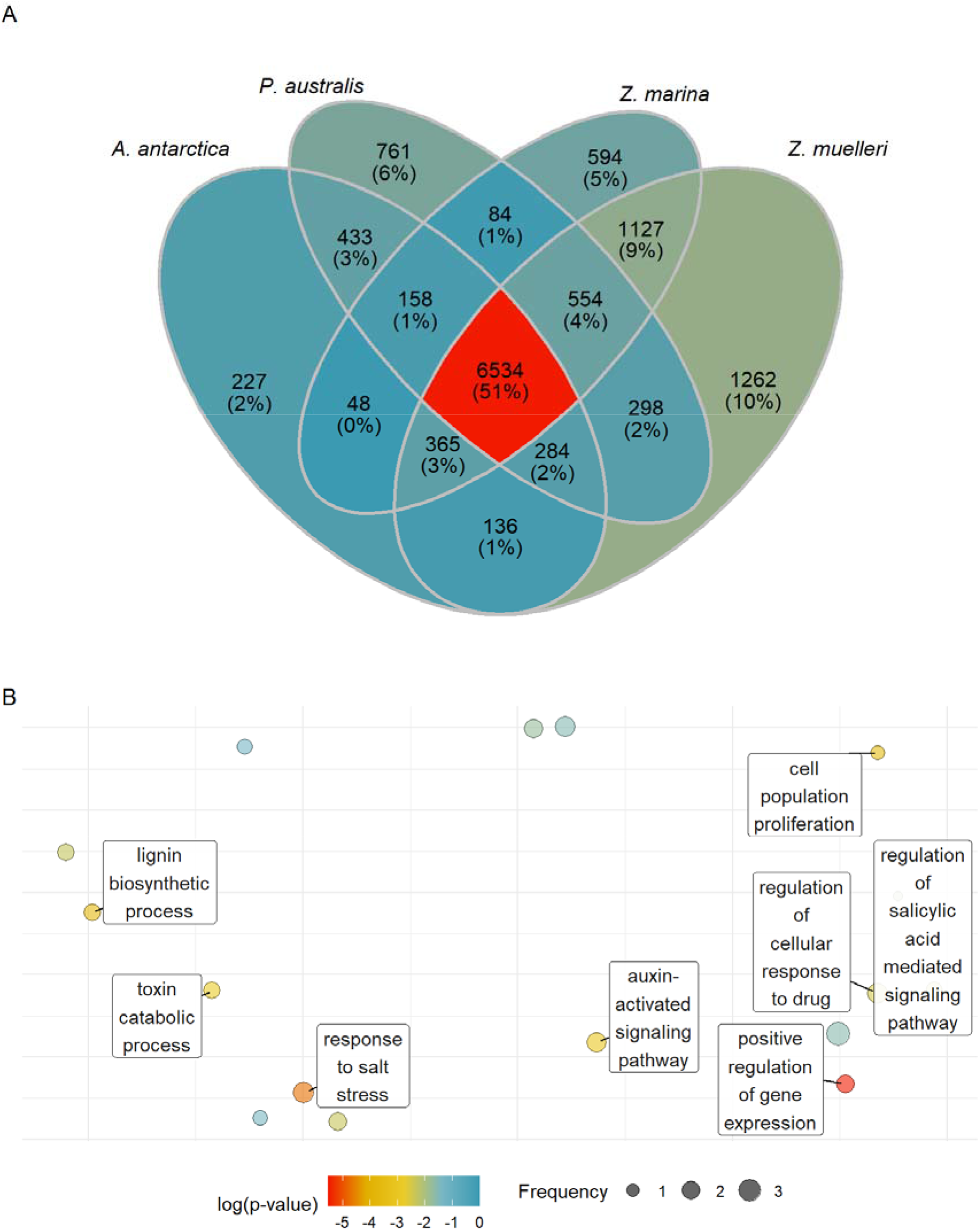
A) Shared and unique orthogroups the four seagrasses showing that 51% of all orthogroups are shared across four species. B) Enriched Gene Ontology (GO)-terms in genes shared by all four seagrasses (intersection), compared with all seagrass GO-terms. Presence of genes was determined by orthogroup membership. The GO-terms are grouped by semantic similarity using REVIGO (Supek et al., 2011). Size of terms is determined by how often similar terms appear in the GO-enrichment output. Dots are coloured by the log of GO-term enrichment adjusted p-values as determined by topGO, GO-terms with an adjusted p-value above 0.05 were removed (Alexa and Rahnenführer, 2009). The eight GO-terms with the lowest adjusted p-values are labelled.

We then compared Gene Ontology (GO)-terms enriched across duckweeds, microalgae, seagrasses, and terrestrial plants, and then across the four seagrasses. Enriched GO-terms present in terrestrial plants but lost in the duckweeds, microalgae, and seagrasses included cell wall biogenesis-related GO-terms such as ‘plant-type cell wall loosening’ (GO:0009828); ‘defense response’ (GO:0006952); ‘terpene biosynthesis’ (GO:0051762) and various morphology-related terms such as ‘meristem development’ (GO:0048509), ‘root meristem growth’ (GO:0010082), and ‘root development’ (GO:2000280) (Figure 4B). The GO-terms ‘RNA-dependent DNA biosynthetic process’ (GO:0006278) and ‘transposition, RNA-mediated’ (GO:0032197’) were enriched in seagrasses. GO-terms enriched in all seagrasses included ‘response to salt stress’ (GO:0009651) and ‘positive regulation of gene expression’ (GO:0010628).

Genes present in duckweeds and microalgae but lost in seagrasses and terrestrial plants were enriched for GO-terms such as ‘response to ethylene’ (GO:0009723), ‘floral whorl development’ (GO:0048438), or ‘wax biosynthesis’ (GO:0010025). Conversely, genes present in seagrasses but lost in duckweeds, microalgae, and terrestrial plants were enriched for GO-terms including ‘proteolysis involved in cellular protein catabolic process’ (GO:0051603) and ‘RNA-dependent DNA biosynthetic process’ (GO:0006278).

Lastly, we compared genes lost in the four seagrass species to determine which processes may be species-specific adaptations instead of general adaptations to aquatic life (Figure 5B). Genes present in all four seagrasses are enriched for GO-terms, including ‘response to salt stress’ (GO:0009651) and ‘positive regulation of gene expression’ (GO:0010628). In both *P. australis* and *A. antarctica*, genes lost compared to *Z. marina* and *Z. muelleri* are enriched for GO-term including ‘RNA-dependent DNA biosynthetic process’ (GO:0006278) and ‘RNA modification’ (GO:0009451).

The reduced morphology complexity of *A. antarctica* roots compared with other seagrasses is reflected in lost genes enriched for ‘lateral root branching’ (GO:0080181). *A. antarctica* has uniquely lost genes related to jasmonic acid signalling potentially due to different signalling pathways (‘cellular response to jasmonic acid stimulus’ (GO:0071395), ‘response to jasmonic acid stimulus involved in jasmonic acid and ethylene-dependent systemic resistance’ (GO:0032260), and ‘jasmonic acid metabolic process’ (GO:0009694)).

Only in *Z. marina* are some genes associated with cell wall genesis lost (‘plant-type cell wall organisation or biogenesis’, GO:0071669). *Z. marina* has also lost genes enriched for arsenite and arsenate transport GO-terms (‘arsenite transport’, GO:0015700, ‘arsenate ion transmembrane transport’, GO:1901684), so it may not be able to tolerate arsenic as well as other seagrasses. *Z. muelleri* has been shown to not accumulate arsenic as well as green or red macroalgae (Maher *et al*., 2011; Price *et al*., 2012), while other seagrasses such as *Halophila ovalis* have been shown to be able to accumulate arsenic (Martin *et al*., 2022). These differences in accumulation ability are likely due to losses of arsenite and arsenate transporters in *Zostera spp*.

Compared with the other seagrasses, *Z. muelleri* has lost genes related to light stimuli (‘cellular response to light stimulus’, GO:0071482) and ‘proteolysis involved in cellular protein catabolic process’ (GO:0051603). Contrary to gene losses unique to single seagrass species, substantially fewer GO-terms were uniquely present in the four seagrasses. No uniquely present GO-terms were found in *A. antarctica*, while five GO-terms were unique in *P. australis*. These five GO terms included ‘starch catabolic process’ (GO:0005983) and ‘response to cadmium ion’ (GO:0046686). The largest number of uniquely present GO-terms were found in *Z. marina* (21 GO-terms, including ‘response to heat’ (GO:0009408)) and *Z. muelleri* (15 GO-terms, including ‘positive regulation of wound healing’ (GO:0090303)).

The large genome size of *P. australis* compared with the other genomes is due to the expansion of repetitive elements and not due to whole-genome duplication. We reannotated repetitive elements in the four genomes to determine whether transposon (TE) expansion and loss led to the difference in genome sizes of the four genomes (Figure 6, Supplementary Table 3). We found more LTR-TEs in the *P. australis* genome compared to the other three genomes, with a total LTR-repeat size of 611 Mbp, compared with 46 Mbp, 124 Mbp, and 164 Mbp in *A. antarctica, Z. marina* and *Z. muelleri*, respectively. The small *A. antarctica* genome contained the lowest proportion of repetitive elements across the compared assemblies (47%). In *Z. muelleri*, the cumulative size of DNA repeat lengths was larger by 115 Mbp compared to *Z. marina* (142 Mbp and 27 Mbp in *Z. muelleri* and *Z. marina*, respectively), while LTR repeat content remained similar (164 Mbp and 123 Mbp, respectively), most likely an outcome of whole-genome duplication with subsequent loss of LTR-TEs. Similar mechanisms of genome size reduction due to LTR-TE loss have been observed in terrestrial plants (Devos *et al*., 2002; Vitte and Panaud, 2005), where a whole-genome duplication led to associated identical repeat expansion followed by class-specific loss of repeats.

**Figure 6:**
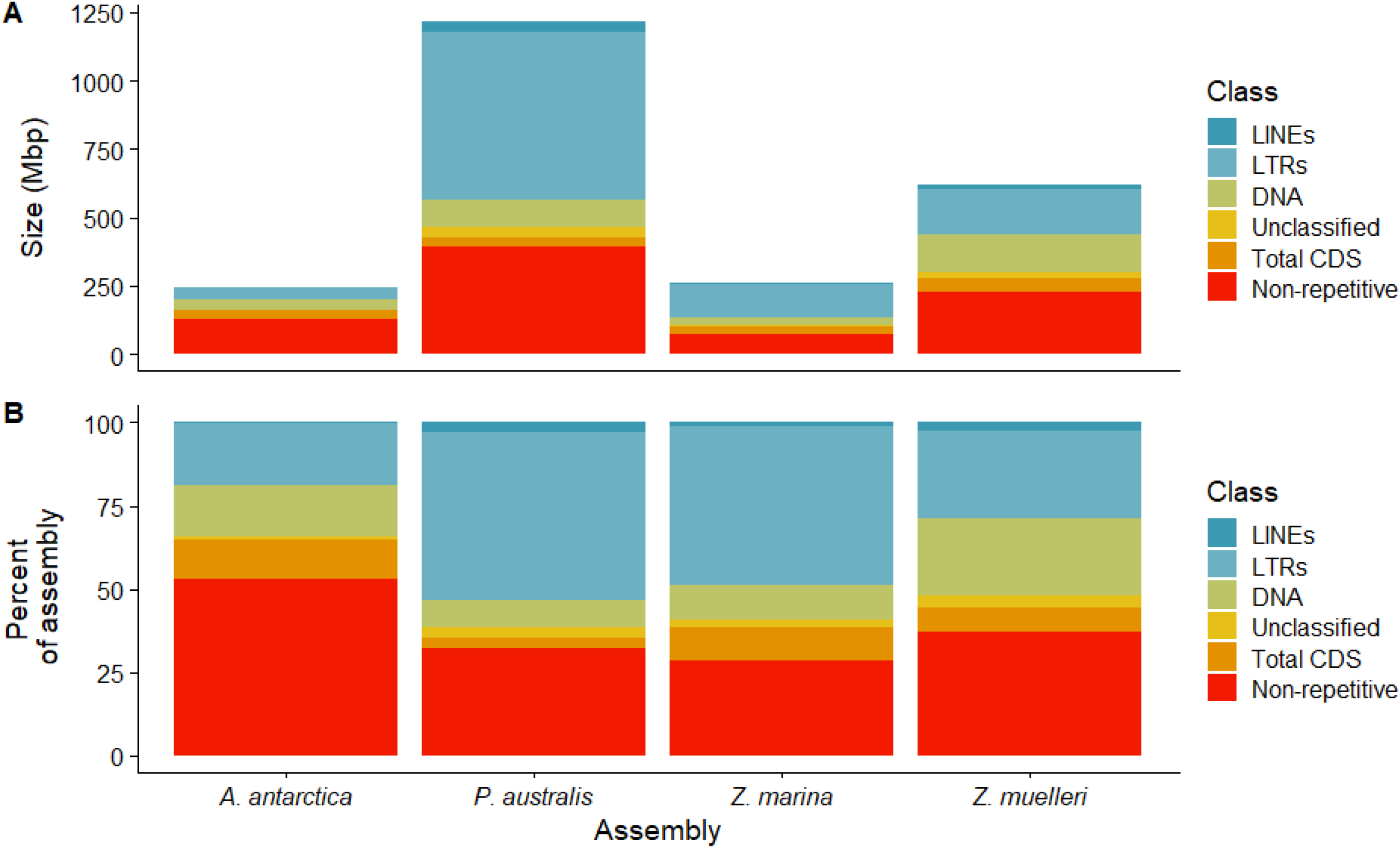
Comparison of repetitive elements in the four assemblies. A) Total size in Mbp of four classes of repetitive elements (LINE, LTR, DNA, unclassified), coding regions, and remaining (non-repetitive) regions. B) Percentage of the total assembly size of four classes of repetitive elements, coding regions, and remaining non-repetitive regions.

We analysed sequence divergence measured by Kimura distance within the four genomes’ TE content (Supplementary Figures 8-11). Only the *Z. marina* genome showed a burst of recent TE activity indicated by low Kimura substitution levels (< 5 Kimura substitution level), with no such recent TE bursts in *Z. muelleri, P. australis* or *A. antarctica*. Interestingly, the TE expansion in the *P. australis* genome is older than in the other seagrasses, with a peak Kimura substitution level of 30 to 35 for DNA-TEs and 40 to 45 for LINE-TEs. Other seagrasses showed younger TE expansion peaks, such as in *A. antarctica* at Kimura distance of 12 and 22 for DNA repeats.

## Discussion

### Two new assemblies for *Amphibolis* and *Posidonia*

Here we presented two new assemblies for *A. antarctica* and *P. australis*. The assemblies are at draft stage, but as they are the first available assemblies for these two families, they form a valuable basis for research into the evolutionary adaptation of plants to life at sea. The genome assembly of *A. antarctica* is the smallest seagrass genome assembly to date, while the *P. australis* genome assembly is the largest available seagrass genome assembly.

There is strong evidence for a whole-genome duplication in *Z. muelleri* (Lee *et al*., 2016), but the larger assembly size of *P. australis* is not due to whole-genome duplication (WGD) as the number of gene models is close to the other non-WGD seagrasses (21,126 gene models compared with 20,450 and 16,176 in *Z. marina* and *A. antarctica*, respectively). A recent study on WGDs in Alismatales also found no evidence for WGD in Posidonia (Chen *et al*., 2022). Rather, the large genome size of *P. australis* is due to greater repeat expansion than *A. antarctica*. In *A. antarctica*, we observed several peaks in Kimura distances between TEs, indicating several rounds of TE expansion, followed by immediate TE shrinkage leading to today’s small *A. antarctica* genome.

One class of repetitive element, MULEs, are entirely missing in *A. antarctica*, indicating that the *Amphibolis* lineage may have diverged from the Posidonia lineage before these transposable elements inserted in seagrass genomes, even though the horizontal transfer of MULEs is widespread in animals (Dupeyron *et al*., 2019). In rice, MULEs have been linked with the rearrangement and amplification of functional gene fragments (Jiang *et al*., 2004), indicating that this mechanism of generating gene diversity may be missing in *A. antarctica*. However, another class of repetitive elements, Helitrons, are also linked with generating gene diversity (Dong *et al*., 2011) and are expanded in the *A. antarctica* genome.

### Convergent adaptation in gene content

As observed in other seagrasses, we observed a reduction of NLR content in *A. antarctica* and *P. australis* (Olsen *et al*., 2016). Similar losses in genes involved in the downstream signalling of NLRs have been observed in aquatic plants. These include convergent loss of the Enhanced Disease Susceptibility 1 (EDS1), Hytoalexin Deficient 4 (PAD4) and Activated Disease Resistance-like 1 (ADR1) disease-downstream signalling pathway in Alismatales (aquatic plants including seagrasses) and Lentibulariaceae (carnivorous plants) (Baggs *et al*., 2020), which is also involved in drought tolerance (Chini *et al*., 2004). This pathway has recently been proposed as a central convergence point for defence signalling cascades (Pruitt *et al*., 2021). As observed in *Z. marina* and *Utricularia gibba (*Baggs *et al*., 2020), we found these three genes were entirely lost in all four seagrasses and three duckweeds, supporting their loss either as adaptation to aquatic life or due to redundancy.

A second resistance pathway found by Baggs *et al*. (2020) to be absent in Alismatales and Lentibulariaceae are the RNLs: *NRG1, NRG1.2, NRG1.3, ADR1, ADR1-L1, ADR1-L2, ADR1-L3*, and *RPW8*. We also found these genes to be absent in all four seagrasses and three duckweeds studied here. Furthermore, we found no RNLs and no protein carrying the RPW8 domain in *A. antarctica* or *P. australis*. The RPW8 domain is present in early land plants and still extant in *Physcomitrella* (Zhong and Cheng, 2016), so these RNLs may have been lost during adaptation to aquatic life or have become redundant. The RNL pathway is also involved in stomatal closure (Chini *et al*., 2004), so it may have been lost as part of the general loss of genes involved in stomatal pathways, as seagrasses do not have stomata. The two new genomes presented here will form the basis for future research into seagrass mechanisms of disease resistance and provide future opportunities to monitor diseases such as wasting disease, which is predicted to worsen with climate change (Graham *et al*., 2021).

We found differences in the ethylene gene content across the studied seagrasses. Unlike Zosteraceae, both ethylene-producing and ethylene-sensing groups of genes are present in *P. australis*. These findings indicate that the ethylene pathway may be functional in *P. australis* as only a few ethylene pathway-related genes are lost and confirm similar findings of a partial presence of ACC synthases and oxidases in Posidoniaceae (Chen *et al*., 2022). Ethylene plays an important role in hypoxia survival and metabolic acclimation during submergence in terrestrial plants (Hartman *et al*., 2021). Seagrasses can experience extended hypoxic stress, particularly at night when low water movement can limit gas exchange, as shown for *Posidonia sinuosa*, a close relative of *P. australis* (Olsen *et al*., 2018). Ethylene may play a similar role in enhancing survival during hypoxia, which may be more prevalent in larger species, such as *Posidonia*, which have greater respiratory demands. However, the role of ethylene in larger seagrasses requires physiological and biochemical confirmation.

In *A. antarctica*, only ethylene-precursor-producing genes ACC synthases are present, while the entire ethylene pathway, including *EIN3* (present in *Z. marina* and *Z. muelleri*) is lost. *A. antarctica* may use ACC to feed bacteria in a mutualistic relationship as has been proposed in terrestrial plants, where many ACC deaminases (ACCD) have been discovered in rhizomes in the presence of plant-mutualistic bacteria (Singh *et al*., 2015). ACC deaminases in bacteria break down ACC into ammonia and α-ketobutyrate, which are then further metabolised. Such relationships have been observed in *Zostera* marina and Z. japonica (Crump *et al*., 2018), but the microbiome of *A. antarctica* has not been studied, and no links between ACC and ACCD are known in seagrasses. The interaction between ACC synthases and mutualistic bacteria in *A. antarctica* will be the focus of future studies. A summary of our model of remaining ethylene pathways in seagrasses is presented in Figure 7. Comparing *EIN3* orthologues at the sequence level revealed a highly conserved core region with no losses of the ‘Ethylene insensitive 3’ Pfam domain, indicating that all *EIN3* orthologues were functional and involved in similar roles across the plants studied.

**Figure 7:**
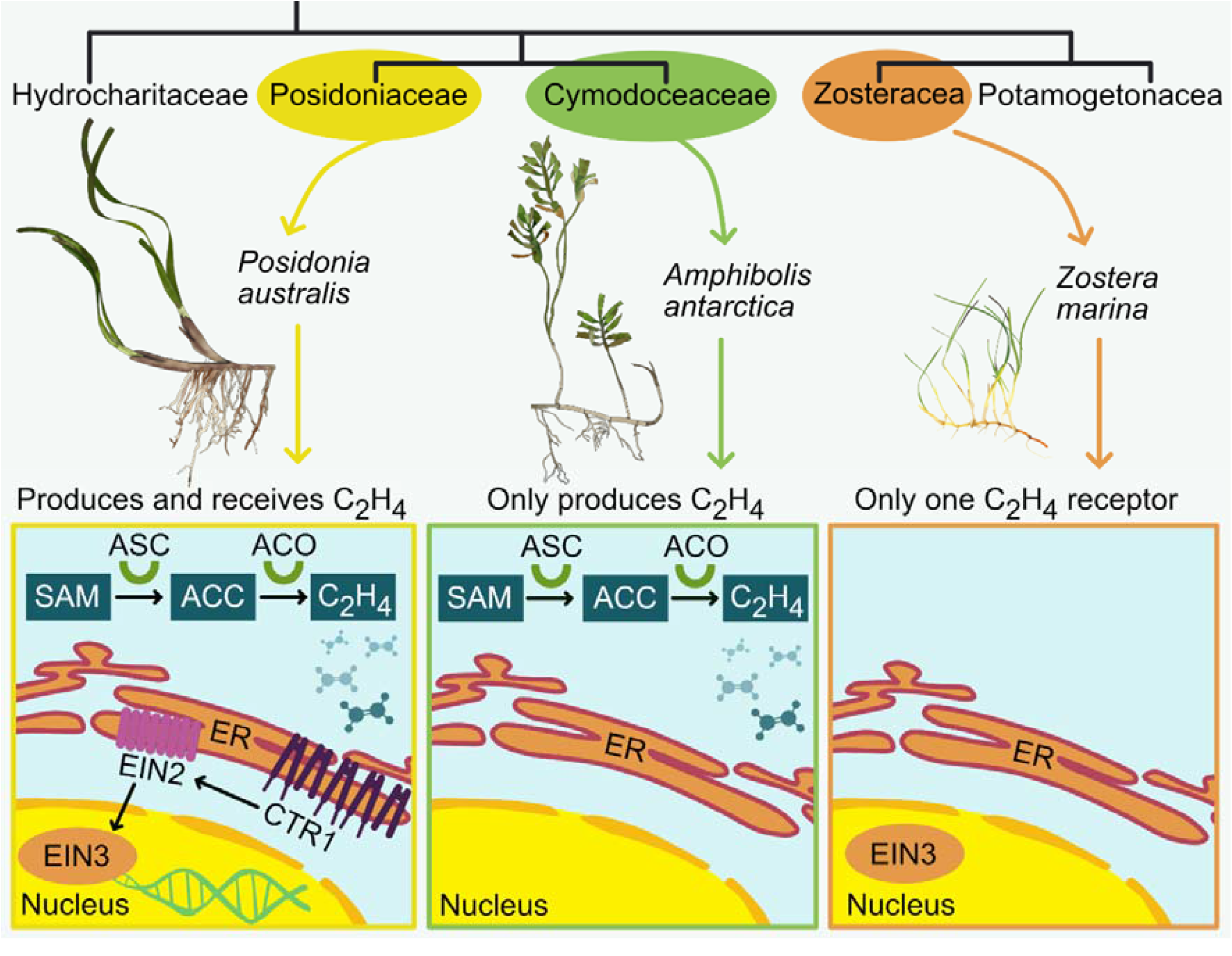
Changes in ethylene pathways in Posidonia australis, Amphibolis antarctica, and *Zostera* marina based on gene presence/absence. P. australis retained the entire ethylene pathway, while A. antarctica can only produce, but not receive ethylene. Z. marina only retained one ethylene receptor.

We found different patterns of *ndh* gene complex losses across the seagrasses studied. The shared presence of *ndhC* and the shared truncated version of *ndhH* and *ndhK* in *A. antarctica* and *P. australis* chloroplast genomes indicates that the *ndh* gene complex is lost in a similar order, with *ndhC* being the last gene to be lost, even though *ndhC, ndhK*, and *ndhJ* are co-located in the same *ndhC* operon and may be expected to be lost together (Meurer *et al*., 1996). Similarly, *ndhA, ndhD, ndhE*, and *ndhG-I* are co-located in the same operon with *psaC*, yet *psaC* is highly conserved and present in all seagrass chloroplast annotations, while most of these surrounding *ndh* genes are lost. This supports the central role of *psaC* in plant survival as it is retained across all four seagrass families.

Four nuclear *ndh*-complex-related genes are present in all four seagrasses, including *PnsL5*. *PnsL5* (also known as *AtCYP20-2*) may not be a subunit of *ndh*, as *A. thaliana* mutants lacking *PnsL5* show unmodified *ndh* activity (Sirpiö *et al*., 2009), so it fulfils an as yet unidentified, but important role causing its retention.

The *PGR5/PRGL1* pathway is the main PS1 cyclic electron transport in angiosperms, along with the chloroplast *ndh*-complex dependent pathway (Shikanai, 2014). We found *PGR5* and *PRGL1* genes to be conserved in the four seagrass genomes. The genes PGR5 and PRGL1A are important for cyclic electron transport and in protecting photosystem I and II (Ma *et al*., 2021a; Munekage *et al*., 2002). This pathway is especially important in protecting photosystems under fluctuating light conditions (Yamori *et al*., 2016), a common feature in aquatic environments. Therefore, conservation of this pathway may have been preferred to PS1 cyclic electron transport functions provided by a loss of the *ndh* pathway.

The fourth *ndh*-complex-related gene conserved across seagrasses is *CRR2. CRR2* encodes for a Pentatricopeptide repeat (PPR) protein (Ruwe *et al*., 2019) which is essential for the expression of n*dhB* (Hashimoto *et al*., 2003). Since *ndhB* is lost in all seagrasses still carrying *CRR2, ndhB* may not be the only binding target of *CRR2*. Further studies will focus on identifying additional binding targets for *CRR2*.

We found all stomata generation-related genes except SCRM2 were lost as seagrasses have generally lost stomata in adaptation to underwater life. Of the stomata-related genes, we found only *SCRM2* to be conserved, with about 50% of its sequence covered by reads of all species and orthologues found in all orthogroups, which confirms the findings of Chen *et al*. (2022). *SCRM2* (*SCREAM2*) encodes a transcription factor typically involved in all stages of stomata generation (Kanaoka *et al*., 2008). However, as a transcription factor, it is also involved in other key processes, especially cold acclimation (Kim *et al*., 2015b), which may have prohibited *SCRM2* from being lost.

We compared cell wall generation-related gene content across the seagrasses. *Z. marina* has lost the cell wall biogenesis-related genes present in *A. antarctica* and *P. australis*, indicating further cell wall-related morphological changes in addition to the general seagrass-related cell wall changes. These changes are confirmed by the lack of lignin in *Zostera c*ell walls, conversely present in the *A. antarctica* and *P. australis* cell walls (Pfeifer *et al*., 2022).

Salinity tolerance is essential for survival in the saline ocean. We observed almost no loss in salinity tolerance genes, but we did identify divergent copies of salinity-pathway-involved genes, especially in *SOS3*. Transgenic *Arabidopsis* plants overexpressing *SOS3* show increased salt tolerance (Yang *et al*., 2009). It is possible to transgenetically introduce SOS copies from halophile plants to increase salinity resistance (Liu *et al*., 2021). The additional diverged copies of *SOS3* in the seagrasses are likely diverged to handle high-salinity conditions, which makes them targets for introduction into other plants. The seagrass copies of *SOS3* contain up to 205 uncharged amino acids at the beginning or end of the protein sequences, which are not present in the terrestrial orthologues. These uncharged regions may represent intrinsically disordered regions. These regions may represent adaptations to marine life as disordered regions have been shown to change conformation in response to salinity (Müller-Späth *et al*., 2010; Wohl *et al*., 2021). Exactly how these conformation changes lead to increased salinity tolerance is currently unknown. The additional *SOS3* copies may help *Zostera spp*. survive in different salinity conditions, but there are no studies comparing seagrass salinity tolerance across salinity gradients. An expanded SOS pathway would explain why *Z. muelleri* can survive as an intertidal and estuarine species, maintaining photosynthesis in as little as 15% salinity of normal seawater (Kerr and Strother, 1985). Another explanation may be sexual reproduction, where freshwater pulses initiate seed germination in *Z. muelleri*, and additional SOS3 copies may assist in responding to freshwater pulses (Stafford-Bell *et al*., 2016).

Terpenoids have a well-known role in biotic and abiotic stress response and are used by many terrestrial plants to defend against herbivores and pathogens, including signalling to beneficial organisms (Pichersky and Raguso, 2018). One example is cotton: when cotton is infested by corn earworm caterpillars, terpenoids are emitted by the plant to attract insects that feed on corn earworm caterpillars (McCall *et al*., 1994). However, many terpenoids have low solubility in water (Martins *et al*., 2017), which implies that insoluble terpenoids cannot be used as a signal underwater or may even be detrimental to seagrasses. High herbivory rates on seagrasses by turtles, dugongs, fish, and invertebrates suggest that non-terpenoid-dependent pathways occur in plant signalling underwater. We found no terpenoid biosynthesis genes in all four seagrasses, except for a single, highly diverged copy in *P. australis*, indicating that seagrasses do not rely on terpenoids for defence signalling.

We compared gene content across terrestrial plants. Most of the GO-terms observed to be lost here were previously reported as lost in seagrasses (Golicz *et al*., 2015; Lee *et al*., 2016; Lee *et al*., 2018; Olsen *et al*., 2016). Genes present in duckweeds and microalgae but lost in seagrasses and terrestrial plants were linked to several GO-terms such as ‘response to ethylene’ (GO:0009723), ‘floral whorl development’ (GO:0048438), or ‘wax biosynthesis’ (GO:0010025) indicating that the loss of these genes was not essential for adaptation to aquatic life.

## Conclusions

We have presented two new seagrass draft genome assemblies for *Amphibolis antarctica* and *Posidonia australis*. These two genome assemblies represent the second lineage of plants, Cymodoceaceae and Posidoniaceae, that have returned to the sea after the previously published genome assemblies for Zosteraceae. We found large differences in genome size, but unlike *Z. muelleri* with a known whole-genome duplication, the size of the large *P. australis* genome is driven by repeat expansion. We compared the four available seagrass genomes and found patterns of gene loss, rather than gene diversification, with many additional species-specific gene losses not directly tied to adaptations to aquatic life. The ethylene pathway is partially present in *A. antarctica* and almost fully present in *P. australis*, which may mean that loss of ethylene pathways may not be associated with adaptations to aquatic life, as previously thought. *A. antarctica* can only ssynthesise ethylene precursors (ACC) and ethylene itself, but all ethylene receptors are lost, which may mean that *A. antarctica* is in a mutualistic relationship with ACC-metabolising bacteria.

Other forms of complex seagrass-bacteria mutualistic relationships have recently been discovered (Tarquinio *et al*., 2019), including the deposition of sugars unavailable to bacteria due to phenolic compounds (Sogin *et al*., 2021) or methane production by archaea living in seagrass meadows (Schorn *et al*., 2022). Future studies will focus on the interaction between the loss of ethylene genes in seagrasses and the role of host microbiomes in providing critical processes for survival (Zilber-Rosenberg and Rosenberg, 2008). Future studies will also include genome assemblies representing the third lineage that has returned to the sea, Hydrocharitaceae, including Najadaceae and Hydrilloideae (*Halophila sp*).

## Methods

### Genome assemblies

Tissue from *P. australis* and *A. antarctica* were collected from Point Peron, Western Australia (−32.2718 S, 115.6898 E) in April 2013, where meadows of both species were growing adjacent. The meristem and youngest innermost leaf were excised from a single plant for *P. australis* and *A. antarctica*. The growth tips were rinsed in sterile water and closely inspected for any visible external contamination. Seven hundred milligrams of tissue were collected and placed in 5ml tubes, flash-frozen in liquid nitrogen, and pulverised using the 2010 Geno/Grinder (SPEX SamplePrep, Metuchen, NJ, USA). A 100 mg of frozen, powdered plant material was transferred to seven 2ml tubes. The DNA was extracted using the Qiagen DNeasy Power Plant mini kit (Qiagen, Hilden, Germany) following the manufacturer’s protocols. The extracted DNA for each of the seven tubes were pooled after elution, and the DNA concentration was quantified with a Qubit fluorometer (Invitrogen, Carlsbad, CA, USA). The quality was confirmed using the LabChip GX Touch (PerkinElmer, Waltham, MA, USA). The extracted DNA was submitted to the Australian Genome Research Facility (AGRF) for library preparation and whole-genome sequencing. Libraries were prepared using the Illumina TruSeq DNA Nano library kit (Illumina, San Diego, CA, USA) according to the manufacturer’s protocol and were sequenced on the Illumina NovaSeq 6000 to generate 100Gb of 150bp paired-end reads. The genomes were assembled using MaSuRCA v4.0.1 (Zimin *et al*., 2013). Transposable elements were annotated using EDTA v1.9.4 (Ou *et al*., 2019) and classified using TESorter v1.3.0 (database: rexdb-plant) (Neumann *et al*., 2019; Zhang *et al*., 2019).

All RNASeq reads were aligned using HISAT2 v2.2.1 (Kim *et al*., 2019). AUGUSTUS v3.4.0 (Stanke *et al*., 2006) and GeneMark-ET v 4.64_(Lomsadze *et al*., 2014) models were trained using the RNASeq alignments via the BRAKER2 pipeline v2.1.6 (Hoff *et al*., 2019). These models along with predicted repeats, were used in MAKER2 3.01.03 (Holt and Yandell, 2011) to produce final gene models.

### Assembly comparison

These gene models were translated into protein sequences which were compared using Orthofinder v2.5.2 (Emms and Kelly, 2019) with protein models of *Spirodela polyrhiza* (Wang *et al*., 2014), *Wolffia australiana* (Park *et al*., 2021), *Ostreococcus lucimarinus* (Palenik *et al*., 2007), *Chlamydomonas reinhardtii* (Merchant *et al*., 2007), *Amborella trichopada* (Amborella Genome Project *et al*., 2013), *Arabidopsis thaliana* (Kaul *et al*., 2000), *Brachipodium distachyon* (Vogel *et al*., 2010), *Oryza sativa* (International Rice Genome Sequencing Project, 2005), *Physcomitrella patens* (Rensing *et al*., 2008), *Popu*l*us trichocarpa* (Tuskan *et al*., 2006), *Selaginella moellendorffii* (Banks *et al*., 2011), *Thellungiella parvula* (Dassanayake *et al*., 2011), Vitis vinifera (Jaillon *et al*., 2007), and Zea mays (Schnable *et al*., 2009).

Seagrass reads were compared using the *Arabidopsis thaliana* annotation using bowtie2 v2.3.5.1 (Langmead and Salzberg, 2012), and per-gene coverage was calculated using mosdepth v0.3.2 (Pedersen and Quinlan, 2018) and bedtools v2.30.0 (Quinlan and Hall, 2010).

Chloroplast genomes were assembled using GetOrganelle v1.7.5.0 (Bankevich *et al*., 2012; Jin *et al*., 2020; Langmead and Salzberg, 2012) using the public *A. thaliana* chloroplast assembly NC_000932.1 as reference (Sato *et al*., 1999). Chloroplast genomes were annotated using Chloe (https://chloe.plastid.org/).

### Protein comparison

Pfam protein domains were predicted using Interproscan v5.55-88.0 (Jones *et al*., 2014; Mistry *et al*., 2021). Proteins of the *EIN3* and SOS3 orthogroups were aligned using MUSCLE v3.8.1551 (Edgar, 2004). The best protein model of evolution was chosen using modeltest-ng v0.17 (Darriba *et al*., 2020).The gene-based phylogeny was calculated using that model for 600 bootstraps using raxml-ng v0.9.0 (Kozlov *et al*., 2019). The resulting phylogenies were drawn using ggmsa v1.1.5 (Yu *et al*., 2020) and ggtree v3.0.4 (Yu *et al*., 2017). Multiple sequence alignments were visualised using JalView v2.11 (Waterhouse *et al*., 2009). GO-annotation was carried out using PANNZER 2 (Medlar *et al*., 2018). GO-enrichment was carried out using topGO v2.44.0 (Alexa and Rahnenführer, 2009) and rrvgo v1.4.4 (Sayols, 2020). All associated GO-terms were used as backgrounds for comparisons involving all plants, while for comparisons involving only seagrass-related GO-terms were used as background.

All other visualisations were generated using R v4.1.0 (R Core Team, 2020) using tidyverse v1.3.1 (Wickham *et al*., 2019) and ggplot2 v3.3.5 (Wickham, 2016). All analyses in this manuscript are fully reproducible using workflowr v1.6.2 (Blischak *et al*., 2019) and are available at https://philippbayer.github.io/Amphibolis_Posidonia_Comparison/.

## Acknowledgements

This work is funded by the Australia Research Council (Projects DP180100668, DP1601004497, DP200100762, DP210101932, and DE210100398). This work was supported by resources provided by the Pawsey Supercomputing Centre with funding from the Australian Government and the Government of Western Australia.

## Author contributions

PEB carried out bioinformatics analysis, and PEB and MWF co-wrote the manuscript. MWF, BCM, EAS, GAK sampled seagrasses, MWF, BCM, JP, ASE, JB, and DE contributed to genome sequencing and genomics analyses. JB and DE assisted in writing the manuscript. All authors edited the manuscript.

## Data availability

Raw read data generated for this study have been deposited in SRA under BioProject PRJNA835502. The *A. antarctica* and *P. australis* genome assemblies were deposited at GenBank under IDJAMKPZ000000000 and JAMKQA000000000.1, respectively. All genome sequences and annotations generated for this study are available at https://www.appliedbioinformatics.com.au/Edwards/index.php/Seagrasses.

