## Supplementary Figures for "Not all pathways are the same – unique adaptations to submerged environments emerge from comparative seagrass genomics"

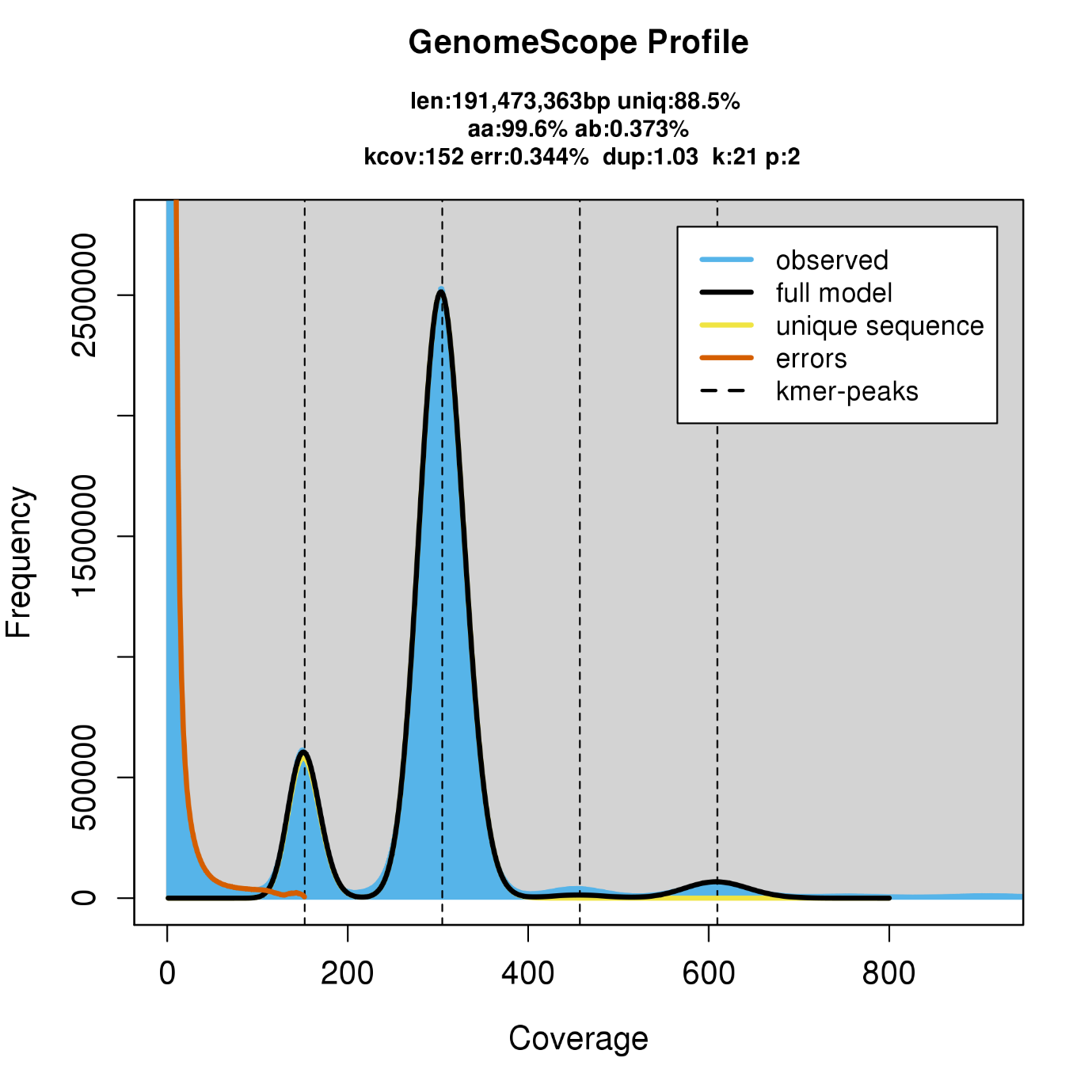

Supplementary Figure 1: GenomeScope v2 genome size estimate model for Amphibolis antarctica

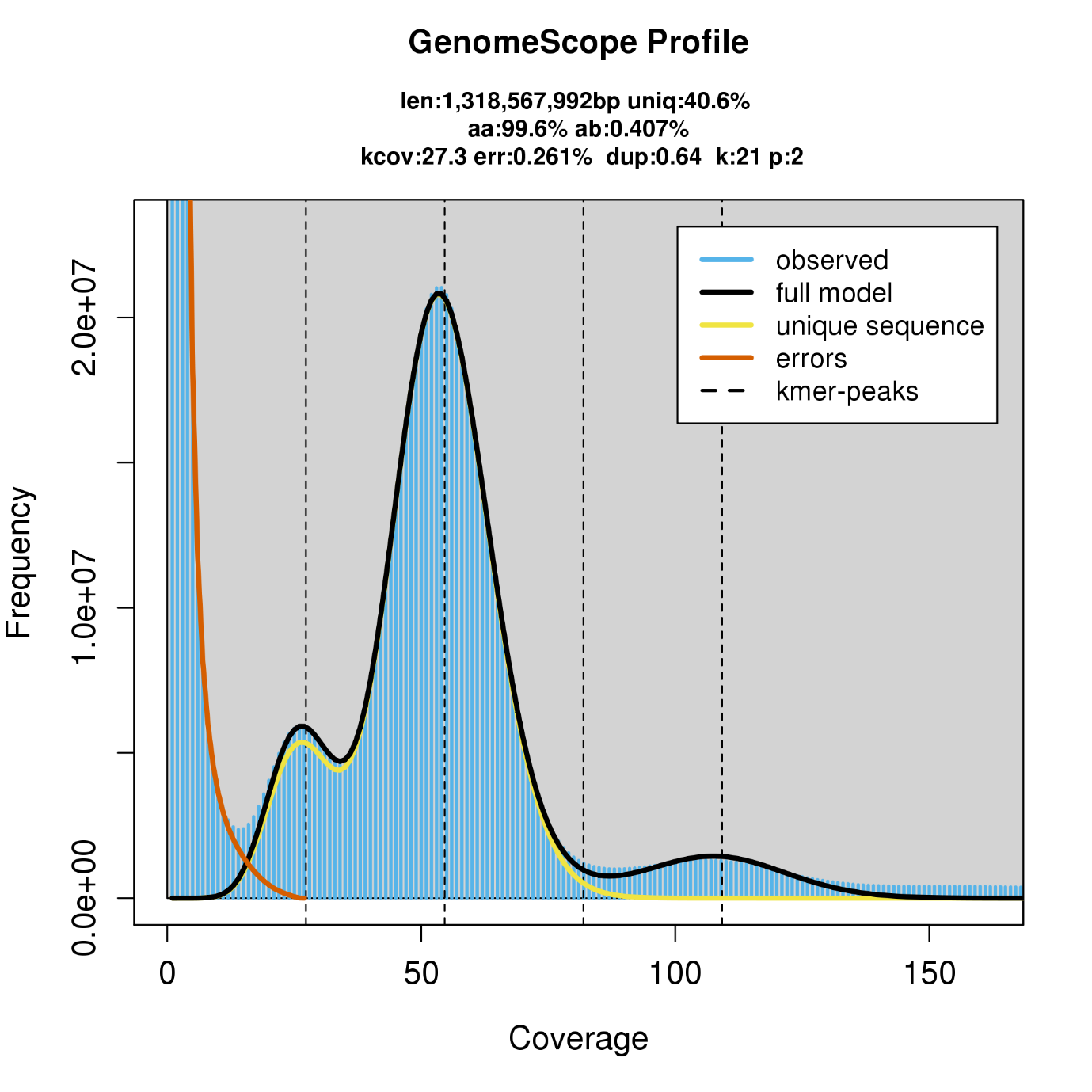

Supplementary Figure 2: GenomeScope v2 genome size estimate model for Posidonia australis

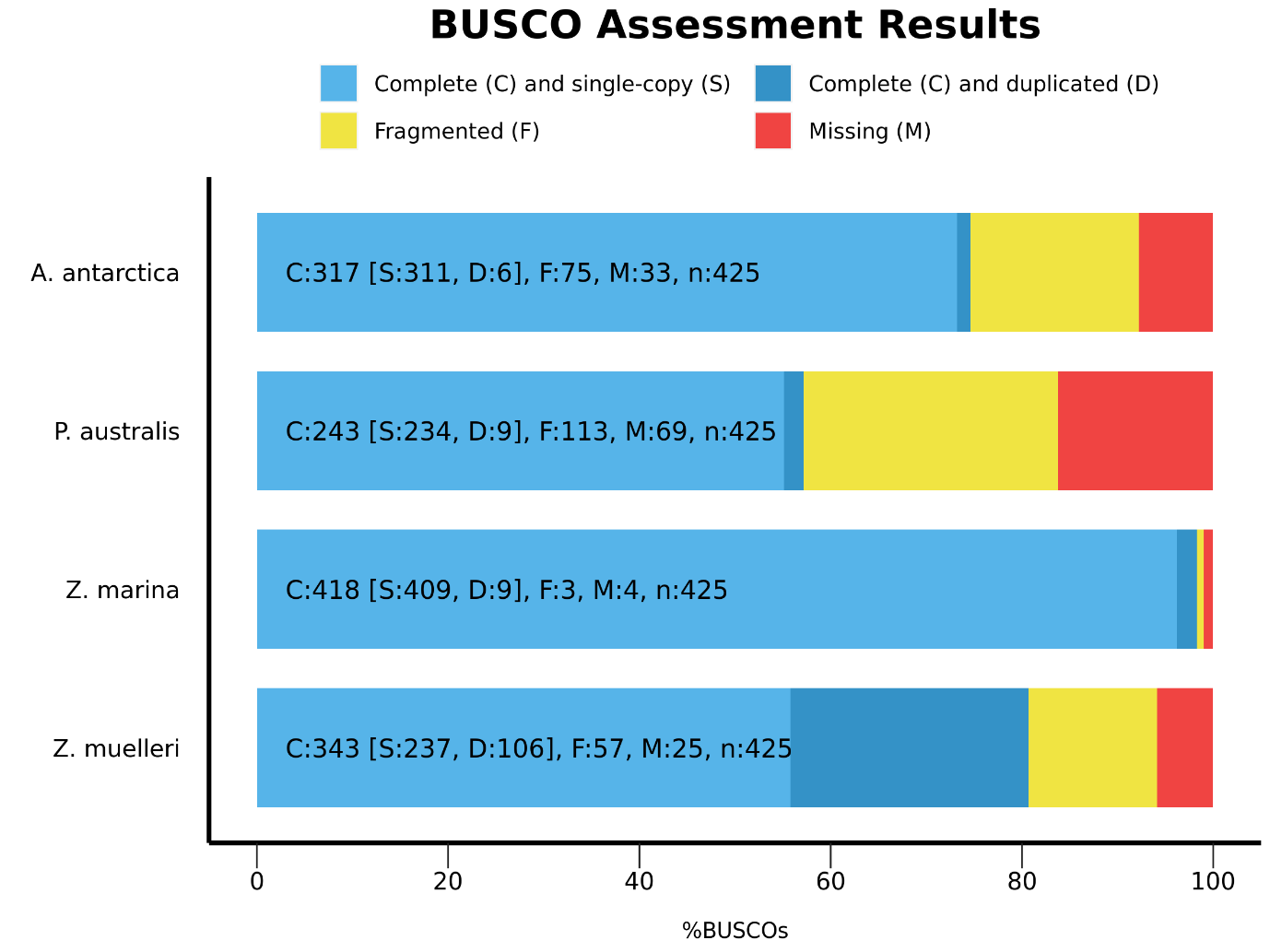

Supplementary Figure 3: BUSCO scores of the four genome annotations using embryophyta v10

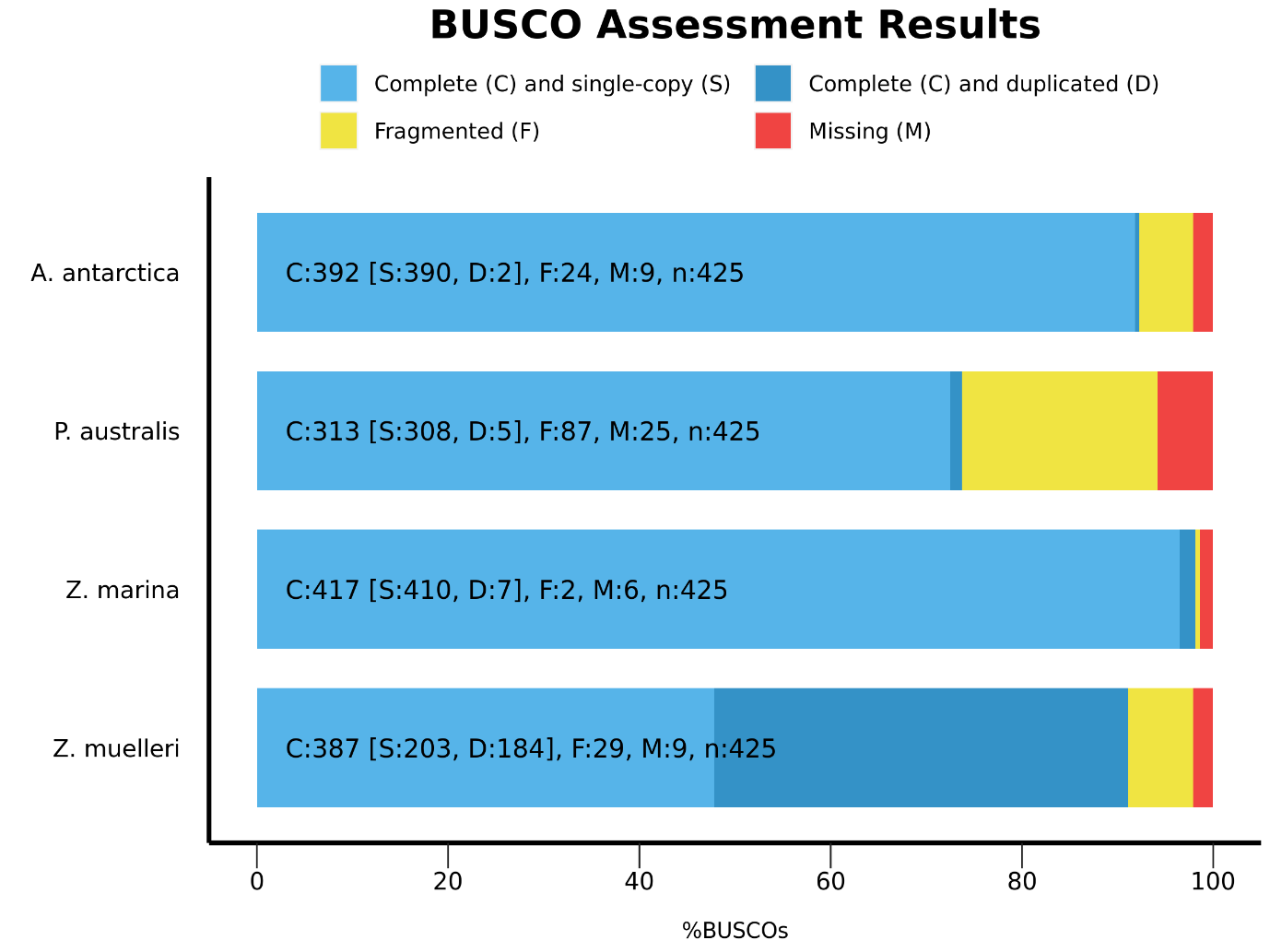

Supplementary Figure 4: BUSCO scores for the four assemblies using embryophyta v10

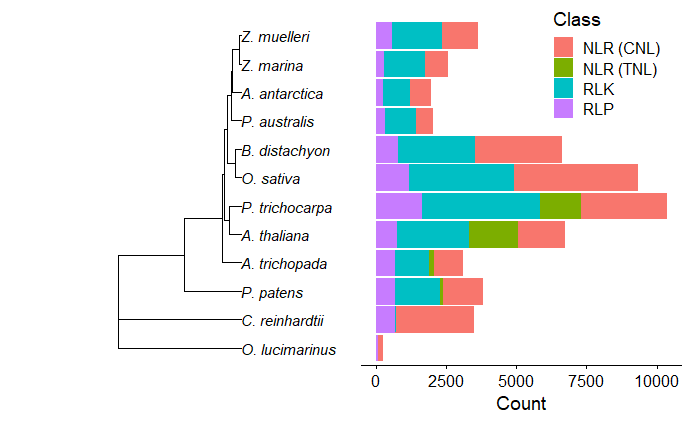

Supplementary Figure 5: Disease resistance gene comparison between the four seagrass genomes and other terrestrial plants and algae showing a reduction in R-genes in all seagrasses compared to terrestrial plants.

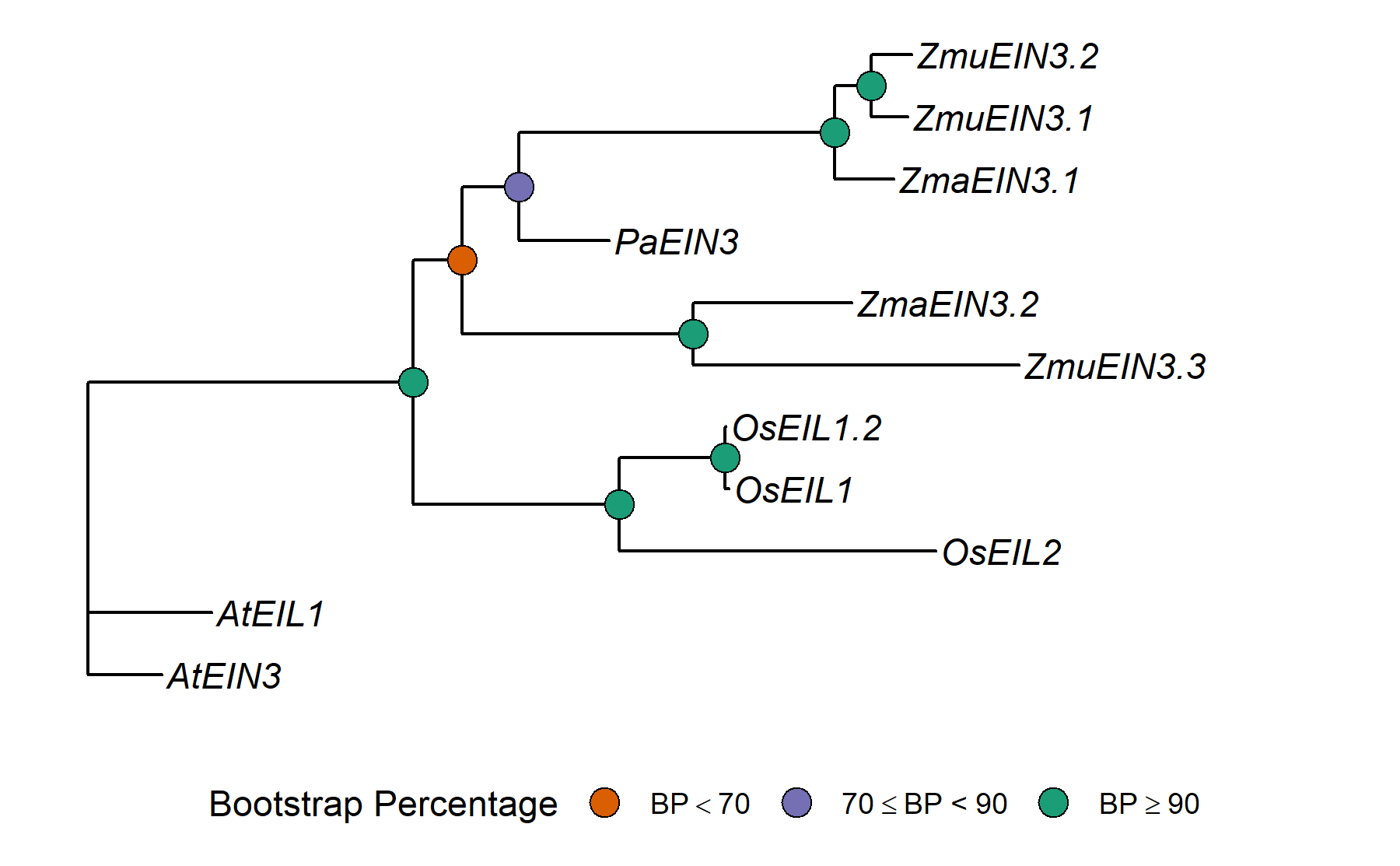

Supplementary Figure 6: Phylogeny of EIN3-included orthogroup of Arabidopsis thaliana (At), Oryza sativa (Os), Posidonia australis (Pa), Zostera marina (Zma) and Z. muelleri (Zmu) genes. Nodes are colored based on bootstrap branch support values from 1000 trees using the model JTT+I+G4+F.

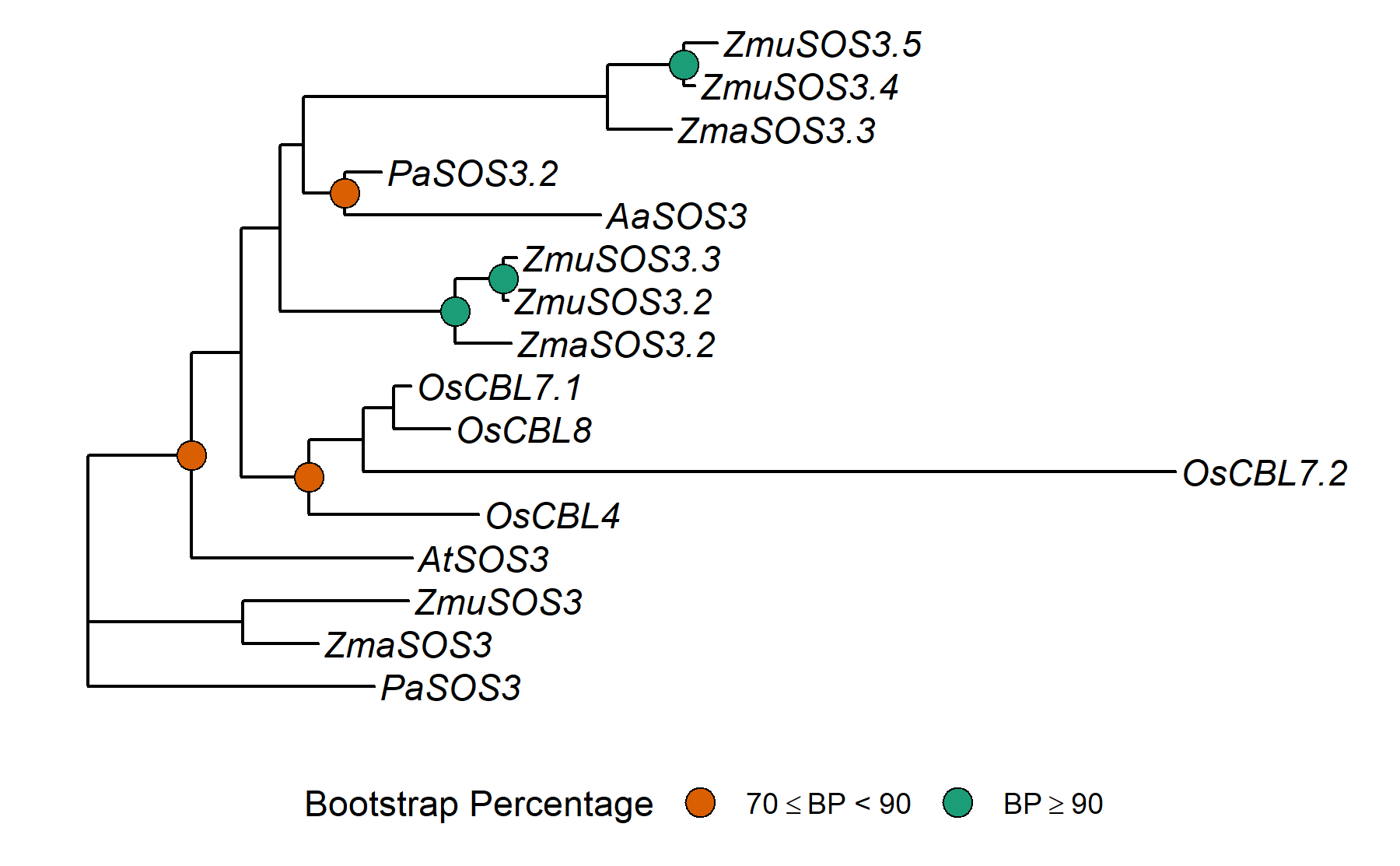

Supplementary Figure 7: Phylogeny of SOS3-included orthogroup of Arabidopsis thaliana (At), Oryza sativa (Os), Amphibolis antarctica (Aa), Posidonia australis (Pa), Zostera marina (Zma) and Z. muelleri (Zmu) genes. Nodes are colored based on bootstrap branch support values from 600 trees using the model JTT-DCMUT+G4.

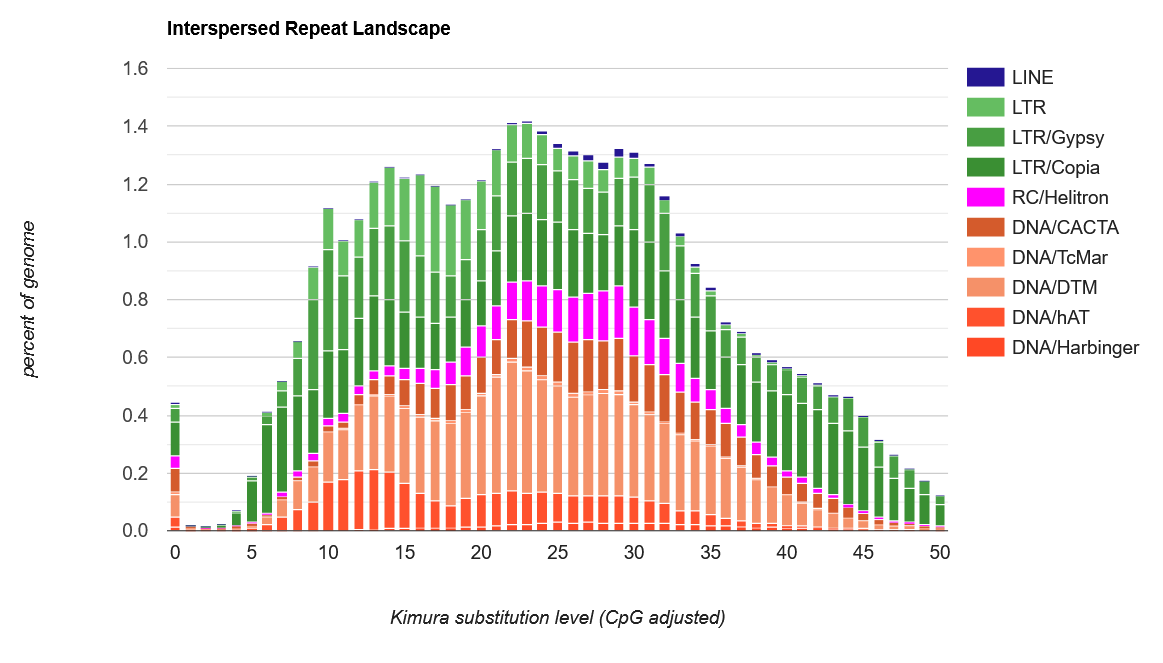

Supplementary Figure 8: Transposable element (TE) accumulation history in the Amphibolis antarctica genome, based on a Kimura distance-based copy divergence analysis of TEs, with Kimura substitution level (CpG adjusted) illustrated on the x-axis, and percentage of the genome on the y-axis.

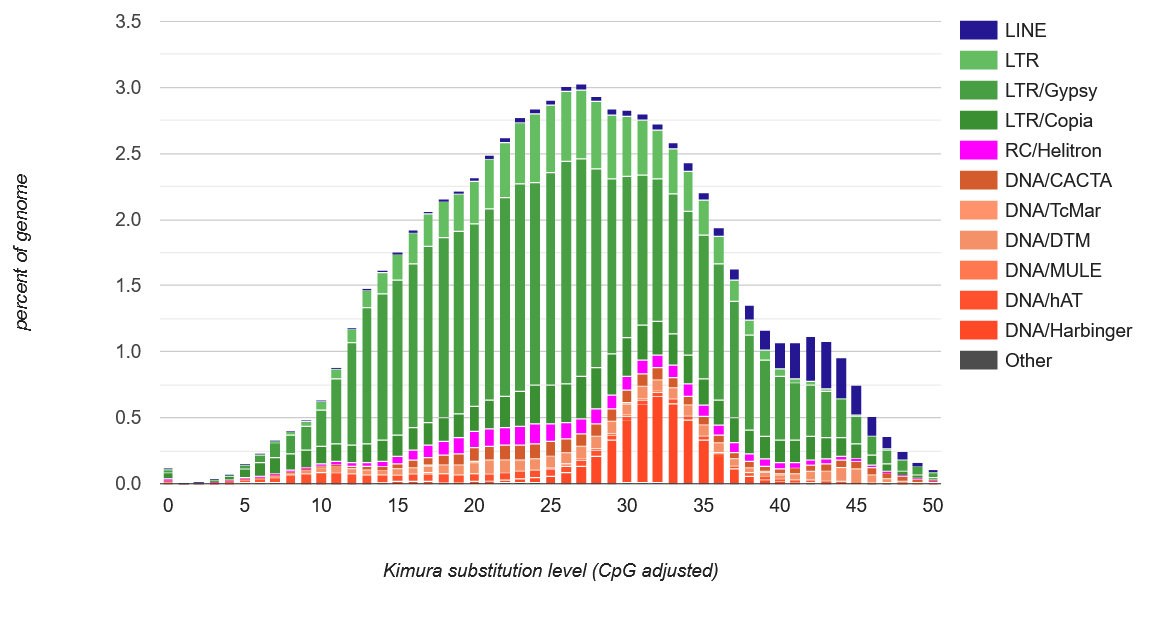

Supplementary Figure 9: Transposable element (TE) accumulation history in the Posidonia australis genome, based on a Kimura distance-based copy divergence analysis of TEs, with Kimura substitution level (CpG adjusted) illustrated on the x-axis, and percentage of the genome on the y-axis.

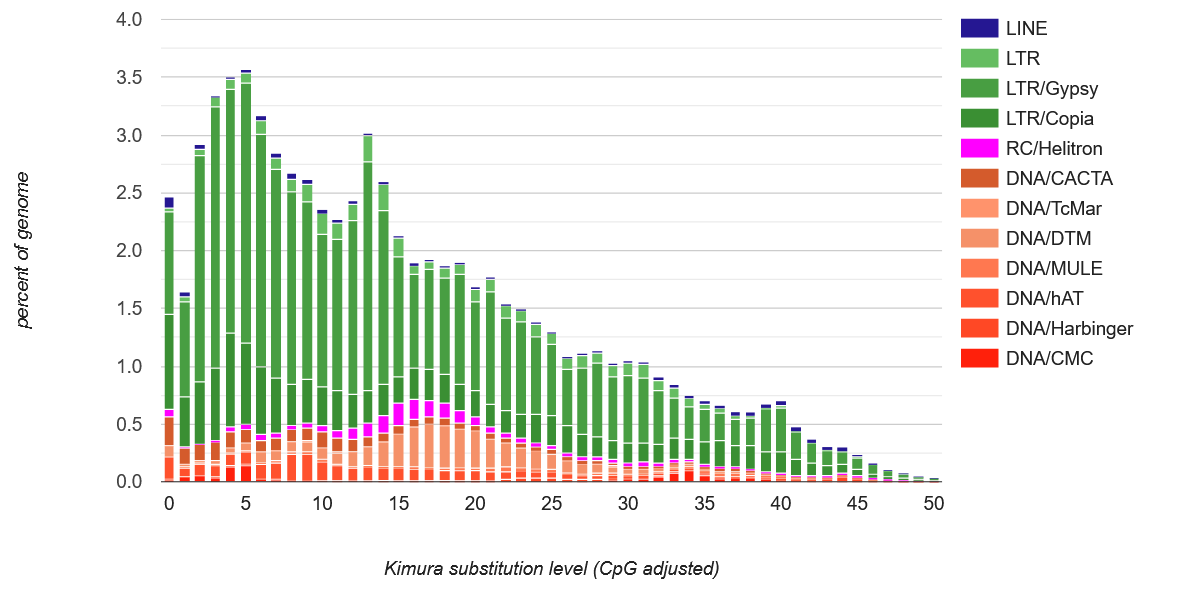

Supplementary Figure 10: Transposable element (TE) accumulation history in the Zostera marina genome, based on a Kimura distance-based copy divergence analysis of TEs, with Kimura substitution level (CpG adjusted) illustrated on the x-axis, and percentage of the genome on the y-axis.

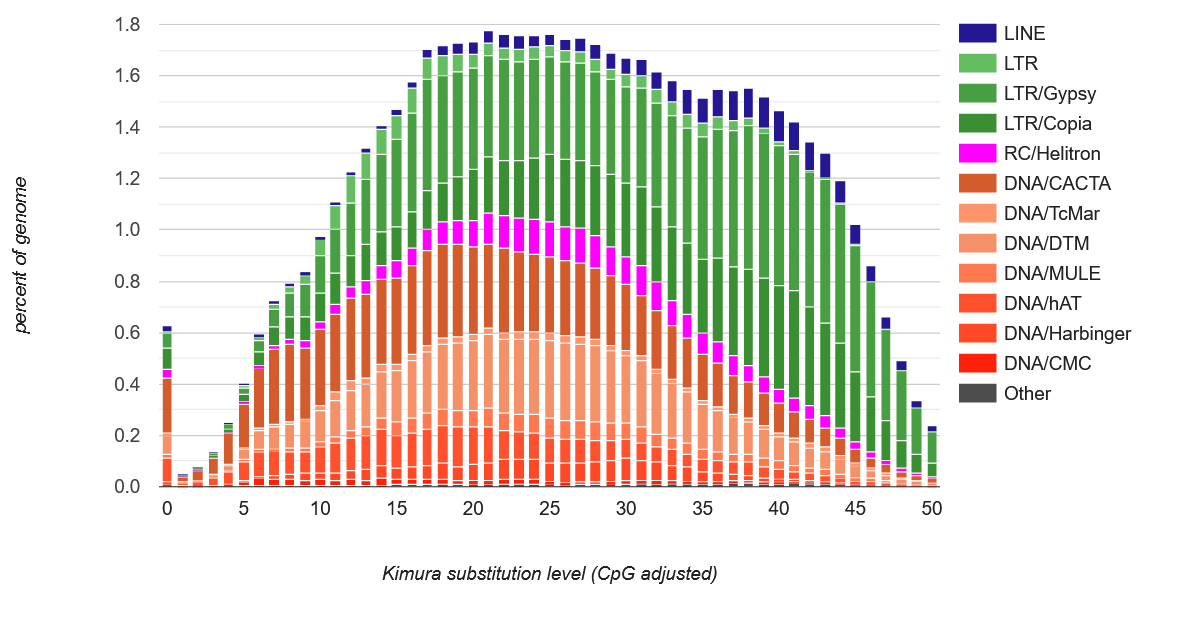

Supplementary Figure 11: Transposable element (TE) accumulation history in the Zostera muelleri genome, based on a Kimura distance-based copy divergence analysis of TEs, with Kimura substitution level (CpG adjusted) illustrated on the x-axis, and percentage of the genome represented on the y-axis.
